# Addition of DMSO to T cell cultures skews differentiation towards a memory phenotype

**DOI:** 10.64898/2026.03.13.711393

**Authors:** Elien De Bousser, Nele Festjens, Evelyn Plets, Leander Meuris, Nico Callewaert

**Author notes:** Two first authors equally contributed (Elien De Bousser, Nele Festjens). Correspondence should be addressed to NF and NC. Author contributions EDB and NF were responsible for the experimental design, data analysis and wrote the manuscript. EDB, NF and EP acquired the data. LM performed the statistical analyses. NC initiated and supervised the project and revised the manuscript. All authors have read and approved the final manuscript.

## Abstract

Dimethyl sulfoxide (DMSO) is a polar aprotic organic solvent that is widely used in biological applications. It is routinely applied as a cryoprotectant for long-term cell freezing as well as to dissolve peptides or drugs for immune cell functional assays. We report on a remarkable impact of low concentrations of DMSO on *in vitro* expansion of the total CD3^+^ pool isolated from human PBMCs as well as on the purified CD8^+^ T cell fraction thereof in the presence of different cytokine combinations typically used for therapeutic T cell expansion. Characterizing survival, proliferation, activation, exhaustion and differentiation, we demonstrate that DMSO at low concentrations substantially skews the differentiation of T cells towards a memory phenotype in a dose-dependent way. This is a desirable outcome for the field of adoptive T cell therapies for cancer, where it has been established that T cells with a memory phenotype exert superior anti-cancer immune responses.

## Introduction

Dimethyl sulfoxide (DMSO) is composed of a highly polar sulfinyl group and two apolar methyl groups. For this reason, as a solvent at high concentrations it is able to solubilize both polar and apolar substances. DMSO is hence extensively used in biological and medical research to dissolve pharmacological substances with therapeutic applications or for immune functional assays^1^. Stock solutions of such compounds dissolved in DMSO need to be diluted to a DMSO concentration that is innocuous to the cell type or organism that is under treatment.

DMSO exerts anti-inflammatory, reactive oxygen species (ROS) scavenger and immunomodulatory effects and therefore exhibits therapeutic potential for the treatment of several inflammatory diseases. In many countries, DMSO is prescribed for a variety of indications including arthritis and interstitial cystitis, and to treat symptoms such as pain, inflammation and intracranial pressure^2^. DMSO is hence a safe, but poorly understood pharmaceutical agent.

Different cell types respond very differently to varying DMSO concentrations and stimulatory conditions^3–7^. Partly because of this, the effects of DMSO on cell biology have been a topic of ongoing debate in scientific literature as reviewed, with reference to clinical applications^8^. DMSO toxicity is mostly explained by its effect on the physical properties of membranes. It can readily penetrate and diffuse into and through hydrophobic barriers such as the plasma membrane, contributing to decreased membrane selectivity and increased cell permeability^9^. DMSO can also act as a ‘chemical chaperone’ in the stabilization of hydrophobic proteins during protein folding^10^. The cytotoxic concentration of DMSO varies largely according to the cell type and experimental conditions adopted^11,12^.

In the course of experimentation on the effect of certain chemical compounds on T cell behavior, where DMSO was tested as a vehicle control, we serendipitously observed that DMSO on its own appeared to have a strong memory phenotype-promoting effect on *in vitro* cultured CD8^+^ T cells. Following these initial observations, we undertook systematic experimentation to better understand the effect of low concentrations (0.1-1.2% v/v) of DMSO on human CD8^+^ T cell proliferation and differentiation characteristics. We also validated that the observed memory differentiation-promoting effect of DMSO extends to total CD3^+^ T cell pool cultures, the cell preparation typically used in the manufacturing of CAR T cells.

## Results

### Impact of low concentrations of DMSO on cultured human CD8^+^ T cell survival, proliferation and activation

CD8^+^ T cells were purified from healthy Red Cross blood donor buffy coats and incubated with human T cell-activator CD3/CD28 Dynabeads for T cell expansion and activation, in the presence of different cytokine combinations (IL-2 versus IL-7/IL-15) and increasing concentrations of DMSO. In order to evaluate the effect of DMSO on the proliferation of T cells in culture, cells were labeled with Tag-it Violet™ proliferation- and cell tracking dye. Flow cytometry analysis was performed on multiple days following start-up until day 11 in culture in order to evaluate T cell activation and differentiation.

The presence of low concentrations of up to 1.2% DMSO in CD8^+^ human T cell cultures does not have a negative impact on survival of the cells, cultured in the presence of either IL-2 (Figure 1.A) or IL-7/IL-15 (Figure 1.B). When cells are cultured in the presence of IL-7 and IL-15 for a prolonged period of time (11 days, Figure 1.B), an inhibitory effect of DMSO on T cell proliferation is observed when 0.6% DMSO or more is added to the cell culture. However, when cells are cultured in the presence of IL-2 (Figure 1.A), a concentration of 0.6% is still tolerated and only the 1.2% DMSO condition shows a marked decrease in T cell proliferation. A concentration of 2.4% DMSO completely inhibits proliferation of human CD8^+^ T cells and results in 0% surviving cells beyond 7 days of culture (Supplementary Figure 2), prompting us to use 1.2% DMSO as highest concentration in further experiments as an easily implemented intervention that mildly slows down CD8^+^ T cell proliferation, while retaining the same viability of the resulting cell preparation.

**Figure 1.**
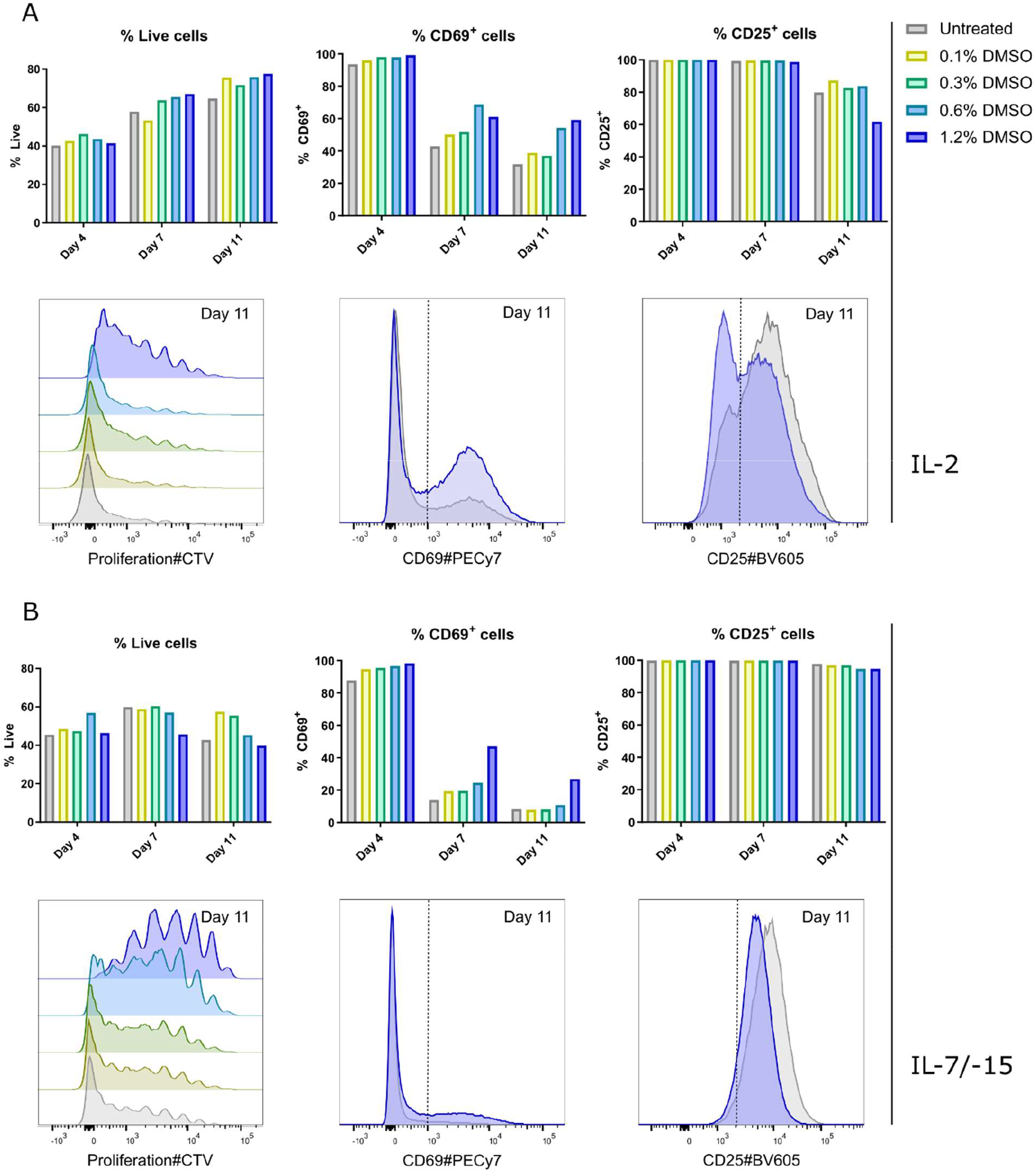
The effect of increasing DMSO concentrations on human CD8^+^ T cell viability, activation and proliferation. Purified human CD8^+^ T cells were stimulated for 4 days and cultured in the presence of IL-2 (A) or IL-7 and IL-15 (B) and increasing concentrations of DMSO (0.1% - 1.2%). Viability, proliferation and expression of the activation markers CD69 and CD25 were analyzed by flow cytometry. Representative histograms are shown for results obtained at day 11 comparing untreated (grey) CD8^+^ T cells with CD8^+^ T cells cultured in the presence of 1.2% DMSO (blue). Representative results are shown from independent experiments performed on T cells isolated from two different blood donors.

The activation markers CD69 (early activation marker) and CD25 (late activation marker) are highly expressed on human CD8^+^ cells freshly activated with CD3/CD28 coated Dynabeads (cells at the end of the 4-day stimulation). The fraction of CD69 positive cells increases with increasing concentrations of DMSO, while the fraction of CD25 positive cells stays similar with increasing concentrations of DMSO, with a minor decrease seen in the 1,2% DMSO condition at day 11 (Figure 1).

### Expression of the negative immune regulator TIM-3 is downregulated on CD8^+^ T cells in the presence of increasing DMSO concentrations

We measured markers that are functionally involved in mediating T cell dysfunction and exhaustion, of which programmed cell death-1 (PD-1), cytotoxic T lymphocyte antigen-4 (CTLA-4) and mucin domain-containing protein 3 (TIM-3). We have analyzed the expression of these exhaustion markers on CD8^+^ T cells in culture in the presence of increasing concentrations of DMSO (Figure 2). As is expected, PD-1 and CTLA-4 are markedly upregulated after anti-CD3/CD28-mediated activation (day 4) and expression levels decrease over time.

**Figure 2.**
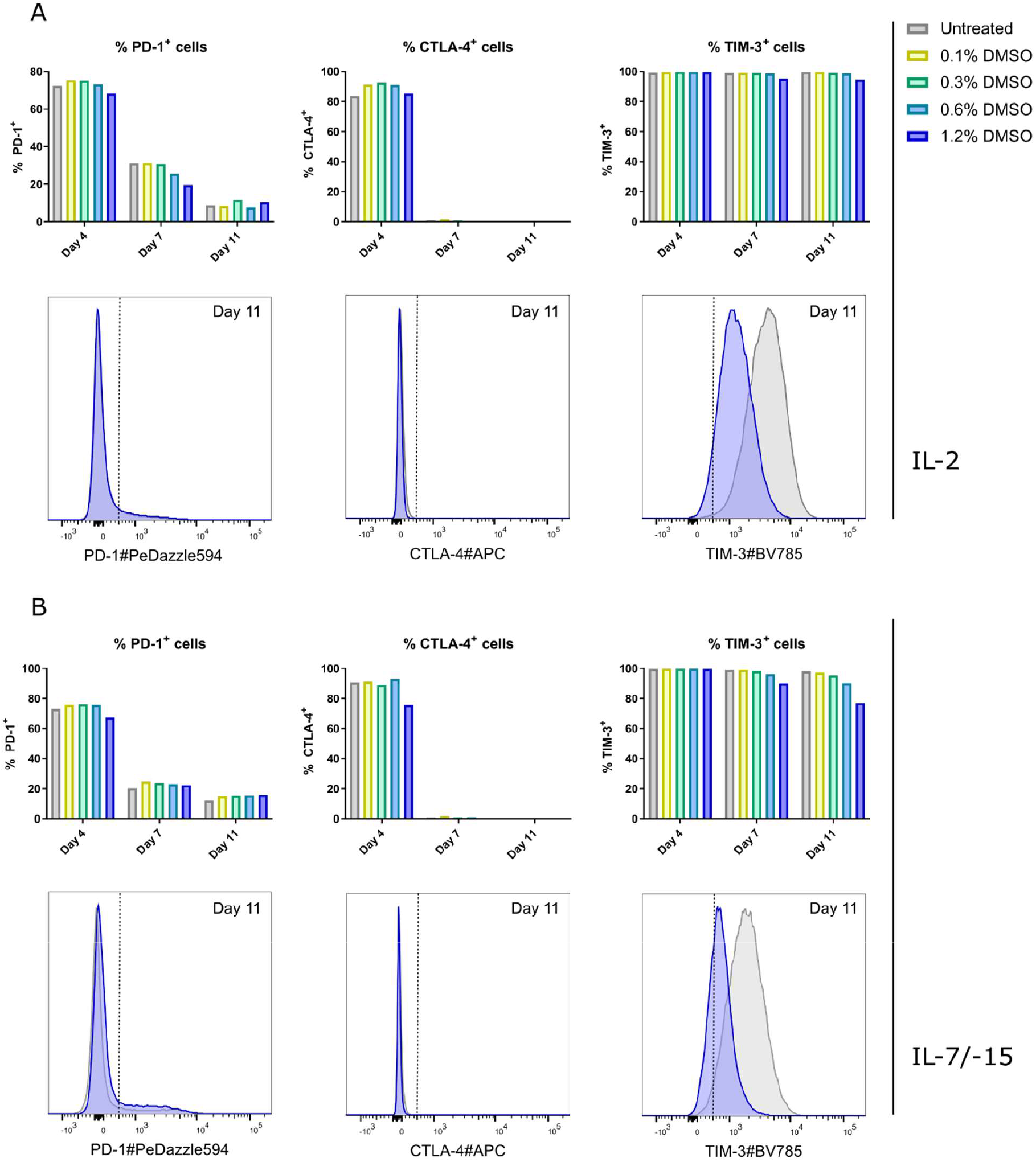
The effect of increasing DMSO concentrations on human CD8^+^ T cell exhaustion. Purified human CD8^+^ T cells were stimulated for 4 days and cultured in the presence of IL-2 (A) or IL-7 and IL-15 (B) and increasing concentrations of DMSO (0.1% - 1.2%). T cell exhaustion characterized by expression of PD-1, CTLA-4 and TIM-3 was analyzed by flow cytometry. Representative histograms are shown for results obtained at day 11 comparing untreated (grey) CD8^+^ T cells with CD8^+^ T cells cultured in the presence of 1.2% DMSO (blue). Representative results are shown from two independent experiments performed on T cells isolated from different blood donors.

The expression of TIM-3 remains high during the entire culturing period of human CD8^+^ T cells in either cytokine condition. This is in line with previous observations showing that TIM-3 is induced by the common γ-chain cytokines IL-2, IL-7, IL-15 and IL-21 in an antigen-independent manner and is upregulated on primary T cells in response to T cell receptor and CD28 stimulation^13^. However, we see a marked decrease in the expression level of TIM-3 in the presence of increasing concentrations of DMSO (Figure 2.B). The TIM-3 expression levels mostly remain above the gating threshold for TIM-3 positivity, but the strong reduction in expression level of this TCR-signaling inhibitory protein could indicate a reduced intensity of TCR signaling upon DMSO treatment.

### DMSO skews T cell differentiation towards a memory phenotype

Human T cell differentiation subsets were quantified during culturing in the presence of different concentrations of DMSO and identified as naive T_N_ cells (CD45RA^+^CD62L^+^ or CD45RA^+^CD197^+^), effector T_EFF_ cells (CD45RA^+^CD62L^-^ or CD45RA^+^CD197^-^), effector memory T_EM_ cells (CD45RA^-^CD62L^-^ or CD45RA^-^ CD197^-^) and central memory T_CM_ cells (CD45RA^-^CD62L^+^ or CD45RA^-^CD197^+^). Untreated cells mainly have a naive-like phenotype by day 11, while DMSO treated cells shift towards a memory phenotype. This skew encompasses both the T_EM_ and T_CM_ populations in the presence of IL-2 and is statistically significant at a DMSO concentration of 0.6% (T_EM_) and 1.2% (both T_EM_ and T_CM_) (Figure 3). The skew towards a memory phenotype also occurs in the presence of IL-7 and IL-15 (Figure 3). In this case, T cell differentiation is significantly skewed towards a T_CM_ phenotype in the presence of DMSO. The decrease in the proportion of naive-like cells and the increase in the proportion of central memory CD8^+^ T cells was significant from 0.3% and up to 1,2% DMSO (Figure 3). Although the results are statistically significant at day 11, the same trend is already appearing at day 7 (data not shown).

**Figure 3.**
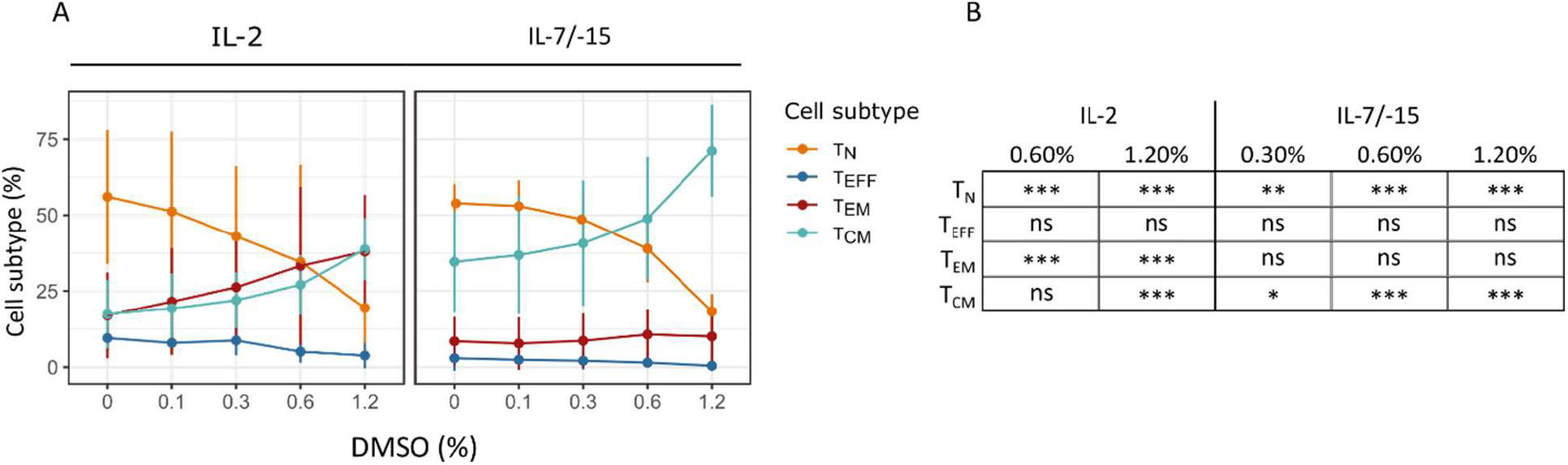
The effect of increasing DMSO concentration on human CD8^+^ T cell differentiation. A. Purified human CD8^+^ T cells were stimulated for 4 days and cultured in the presence of IL-2 (left panel) or IL-7 and IL-15 (right panel) and increasing concentrations of DMSO (0.1% - 1.2%). T cell differentiation at day 11 was analyzed by flow cytometry and subsets are described based on different marker combinations as naive (CD45RA^+^CD197^+^ and CD45RA^+^CD62L^+^), effector (CD45RA^+^CD197^-^ and CD45RA^+^CD62L^-^), effector memory (CD45RA^-^CD197^-^ and CD45RA^-^CD62L^-^) and central memory (CD45RA^-^CD197^+^ and CD45RA^-^CD62L^+^). Results from two independent donors, combined over the different marker combinations for each cell type subset are shown as mean +/− SEM. Statistical analysis (beta regression model) was performed on all cell subtypes comparing each DMSO concentration to the 0% control. Statistically significant differences are indicated in the table in panel B. * *p* < 0.05, ** *p* < 0.01, *\*\*\* p < 0*.*001*

### Evaluation of the impact of DMSO on the CD3^+^ T cell pool used for CAR T cell preparation

Although CD8^+^ T cells are critical for the efficacy of adoptive cell transfer (ACT) due to their cytolytic activity, co-transferred CD4^+^ T cells are known to enhance CD8^+^ T cell-mediated tumor rejection^14^. This is why the total CD3^+^ T cell population is most often used for the production of therapeutic CAR T cells. Several studies have demonstrated that memory T cells, including stem cell memory T_SCM_ and T_CM_ cells show superior persistence and anti-tumor immunity compared with T_EM_ and T_EFF_ cells^15,16^. Unpurified human CD3^+^ T cells stimulated in the presence of IL-12 and cultured in the presence of either IL-2 or IL-7/-15 are also skewed towards a naive-like phenotype. To investigate the effect of DMSO addition on the activation and differentiation of total human CD3^+^ T cells, we decided to evaluate the impact of the DMSO concentration that robustly altered the CD8^+^ T phenotype (1.2%) on the immunophenotype of CD3^+^ T cell (sub)populations at day 13. The latter timepoint was chosen since CD3^+^ T cells engineered *ex vivo* for the study of immune therapeutic applications are generally cultured *in vitro* for a duration of two weeks prior to functional analyses or therapeutic use in the context of CAR T cells.

A DMSO concentration of 1.2% did not impact CD3^+^ T cell survival nor CD4^+^/CD8^+^ subset distribution (Figure 4.A and Supplementary Figure 2). It also does not hamper CD3^+^ proliferation, while this is the case for a 2.4% DMSO addition (Supplementary Figure 2). The frequency of cells that are CD69 positive increases in the presence of DMSO (Figure 4.B), as was seen before (Figure 1). This effect can be seen on both CD4^+^ and CD8^+^ T cell subtypes (Figure 4.B). The effect for CD25^+^ seems to be minor and more pronounced on CD8^+^ T cells (Figure 4.C). Analysis of the exhaustion marker PD1 shows that the proportion of cells expressing PD-1 seems to slightly increase when cells are cultured in the presence of 1.2% DMSO (Figure 4.D), as seen in Figure 2.B.

**Figure 4.**
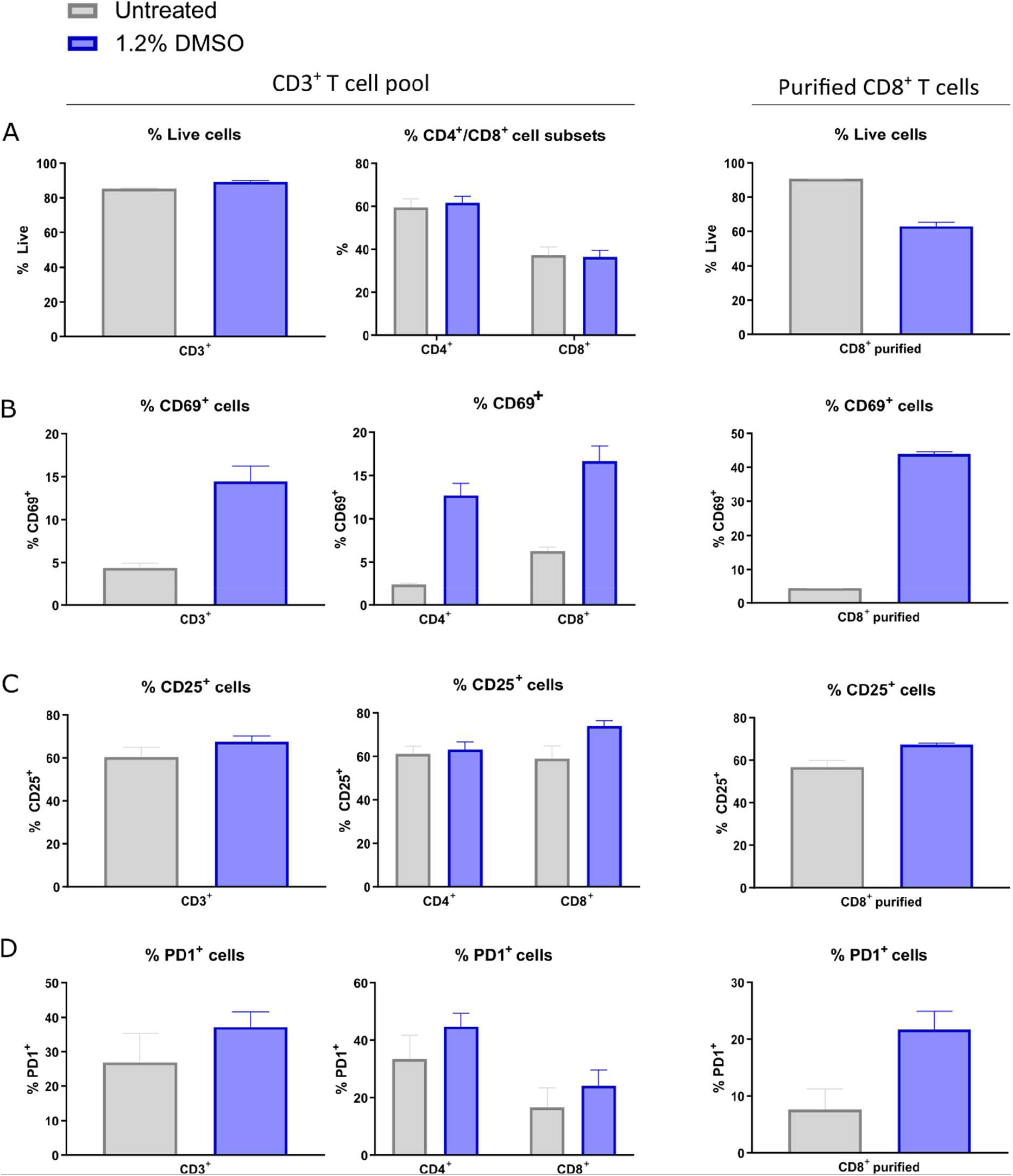
The effect of 1.2% DMSO on human CD3^+^ and CD8^+^ T cell viability and activation. Purified human CD3^+^ T cells and CD8^+^ T cells (from the same donor) were stimulated for 3 days in the presence of IL-12 and cultured in the presence of IL-7 and IL-15 and 1.2% DMSO. Viability (A-left panel) and %CD4^+^/CD8^+^ T cell subtypes (A-middle panel) in the CD3^+^ pool and viability of purified CD8^+^ T cells (A-right panel) were analyzed by flow cytometry. Activation markers CD69 (B) and CD25 (C) and exhaustion marker PD-1 (D) were analyzed by flowcytometry, on either the CD3^+^ pool (left panels) or CD8^+^ purified T cells (right panels) and CD4^+^ and CD8^+^ subtypes in the CD3^+^ pool (middle panels). Representative histograms are shown for results obtained at day 13. Error bars represent standard deviations. The data shown are representative of three independent experiments performed on T cells isolated from three different blood donors.

Importantly, we could demonstrate that in this CD3^+^ T cell expansion protocol, DMSO skews CD3^+^ T cell differentiation towards a central memory phenotype. This skew is more pronounced in the CD4^+^ T cell subpopulation (Figure 5) as compared to the CD8^+^ T cell subpopulation in which only minor changes are observed (Figure 5). In parallel, CD8^+^ T cells were purified from the same donor and cultured in the presence of DMSO. In the latter condition, CD8^+^ T cell differentiation was clearly skewed from naive-like towards a T_CM_ phenotype (Figure 5), confirming earlier results described above.

**Figure 5.**
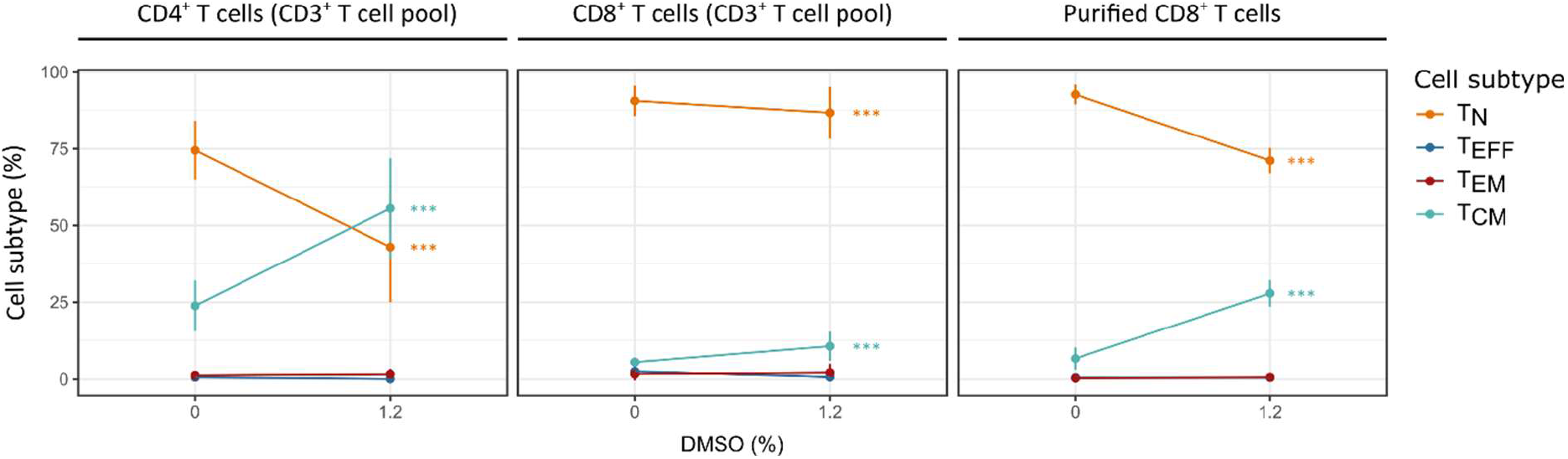
The effect of increasing DMSO concentration on human CD3^+^ and purified CD8^+^ T cell differentiation. Purified human CD3^+^ T cells and human CD8^+^ T cells purified from the same donor were stimulated for 3 days in the presence of IL-12 and further cultivated in the presence of IL-7 and IL-15 without DMSO (0%) and 1.2% of DMSO. Three independent experiments were performed on T cells isolated from three different blood donors. T cell differentiation at day 13 was analyzed by flow cytometry and subsets are described as naive (T_N_, CD45RA^+^CD62L^+^), effector (T_EFF_, CD45RA^+^CD62L^-^), effector memory (T_EM_, CD45RA^-^CD62L^-^) and central memory (T_CM_, CD45RA^-^CD62L^+^) T cells. Results obtained from three independent experiments with T cells isolated from three different blood donors are presented as mean +/− SEM. Statistically significant differences in the fraction of a cell subtype between 0 and 1.2% DMSO are indicated with *** p < 0.001 when significant in the beta regression model.

## Discussion

In this study, we showed that supplementation of T cell cultivation medium with DMSO at low concentrations affects CD8^+^ T cell activation and differentiation *in vitro*. Concentrations up to 1.2% can be used in human CD8^+^ T cultures in the presence of IL-2 or IL-7/-15 without hampering survival. In the presence of IL-2, concentrations up to 0.6% DMSO do not have an impact on proliferation either, which is in contrast to what was previously observed when CD4^+^ T cells are cultured in the presence of low concentrations of DMSO^17^. Importantly, in the typical practice of total CD3^+^ T cell cultivation as is used in CAR T cell manufacturing, concentrations of 1.2% DMSO allowed full viability.

When evaluating the expression of activation markers, we observed that CD69 expression was elevated with increasing DMSO concentrations. It has been described that a small fraction (1-20%) of circulating memory T cells expresses CD69, the latter mediating homing and tissue retention of memory T cells in the bone marrow^18^ and several other organs^19^. We hypothesized that the increased proportion of CD69-expressing cells seen in these initial observations with increased concentration of DMSO, might be indicative of an increase in memory phenotype.

Further, a decrease in TIM-3 expression was observed upon culturing in the presence of DMSO. Most importantly, we demonstrate that addition of DMSO to *in vitro* CD3^+^ and purified CD8^+^ T cell cultures skews the T cell differentiation state towards a memory phenotype.

The assembly of results could be explained by DMSO resulting in some way in a more moderate, but more sustained TCR-downstream signaling during these cultivation protocols, explaining both lower levels of TIM-3 and increase in memory cell formation.

In particular, a decrease in CD45RA expression is observed in the presence of DMSO. CD45 is a tyrosine phosphatase that is alternatively spliced to generate isoforms of different molecular weights which are differentially expressed on T cells depending on their differentiation state. CD45RA^+^ T cells have naive characteristics while CD45RO^+^ (CD45RA^-^) T cells are considered memory T cells because they proliferate in response to recall antigens. Further, a role has been assigned to CD45RA in regulating the cell cycle. CD45RA expression is low during the G0 and G1 stages of the cell cycle^20^. In previous studies, where the effect of DMSO on pluripotent stem cell differentiation was evaluated, it was shown that DMSO increases the time cells spend in the G1 phase of the cell cycle^21^. We therefore hypothesize that the effect of DMSO on the cell cycle of T cells is reflected in a lower expression of CD45RA which is further associated with a memory-like phenotype.

DMSO concentrations ranging from 2.5% to 10% were shown by others to have an *in vitro* anti-inflammatory effect by reducing lymphocyte activation, revealed by reduced proliferation and decreased cytokine production (TNF-α, IFN-γ and IL-2)^12,22^. It the same study, it was shown that treatment with DMSO was not cytotoxic to blood cells when tested at final concentrations up to 1% and only slightly cytotoxic at 2% to 5%^22^. The immunosuppressive activity of DMSO was also demonstrated by Lin et al.^23^ using (0.5 and 1%) DMSO *in vitro*. They showed that addition of DMSO decreases the percentage of IFN-γ-producing T lymphocytes and increases the percentage of Treg cells by enhancing the activation of the STAT5 signaling pathway in the context of autoimmune diabetes. Also in this regard, DMSO was shown to stimulate the production of transforming growth factor-β (TGF-β), an immunosuppressive cytokine, as well as the recruitment of its receptor from intracellular vesicles to the plasma membrane^24^. None of the latter studies documented on DMSO-induced cell death.

We studied the effect of DMSO at much lower concentrations than the high ones used in these studies, which we find to be highly toxic, easily explaining any reduced cellular activity. Indeed, Avelar-Freitas et al. observed that concentrations up to 1% v/v of DMSO did not change IFN-γ and TNF-α production in the cytoplasm of human lymphocytes and neutrophils stimulated with PMA for 4 hours^25^.

Immunotherapy clinical trials based on reinfusion of autologous cells (adoptive cell therapies) often include solvents such as DMSO in their preparation process. For example, the FDA-approved CD19 CAR T cell products Kymriah and Yescarta contain DMSO, used as cryoprotectant. Hence, the molecule clearly is compliant with pharmaceutical use in the setting of adoptive T cell-based therapies. Our observation that either CD3^+^ T cells or purified CD8^+^ T cells can be skewed towards a memory phenotype can be advantageous when preparing T cells for adoptive transfer, since ACT of purified naive, stem cell memory and central memory T cell subsets results in superior persistence and antitumor immunity compared to ACT of populations containing more differentiated T cell subsets such as effector T cells and effector memory T cells^15,16,26^. For instance, it has been shown that deletion of cBAF activity by pharmacological or genetic inhibition of cBAF in CAR T cells could majorly improve their anti-tumor potential *in vivo*, most likely by biased memory differentiation post-ACT^27,28^.

Our data may thus have important implications for future design of cell-based immunotherapies. Simple addition of low, non-cytostatic concentrations of DMSO to T cell cultures for ACT including CAR T cells, might have a significant beneficial impact on their antitumor efficacy *in vivo*, and will thus be subject of our further studies.

## Materials and methods

### Ethical approval

All experiments were approved and performed according to the guidelines of the committee on Medical Ethics of Ghent University, Belgium.

### Human CD8^+^ T cell isolation and culture under DMSO exposure

Leukocyte-enriched buffy coat samples were obtained from healthy donors attending the Red Cross center after informed consent and ethical committee approval (EC2019-1083). Peripheral blood lymphocytes were prepared by a Ficoll-Paque density centrifugation as described in the instruction manual for Leucosep™ (Greiner bio-one). CD8^+^ T cells were isolated by negative selection (MojoSort™ Human CD8^+^ T cell Isolation kit, Biolegend) according to the manufacture’s protocol. Cells were cultured in RPMI 1640 medium (Gibco-BRL) supplemented with L-glutamine (0.03%, Gibco), 0.4 mM sodium pyruvate (Merck Millipore) and 50µM β-mercapto-ethanol (Sigma Aldrich) and 10% heat-inactivated fetal calf serum (FCS), in 48 well plates (Sarstedt) and stimulated with human anti-CD3/CD28 Dynabeads (1 bead:3 cell ratio) (Thermo Fisher Scientific) for 4 days at 37°C. Prior to cell seeding, Dynabeads and bead-bound cells were removed by separation on a magnet (after transfer to an Eppendorf tube) and the supernatant containing the CD8^+^ T cells was collected. Cells were washed twice with PBS before putting them in culture with different cytokine combinations: (1) recombinant human (rh) IL-2 at 50 units/mL (Life Technologies); (2) rhIL-7 at 10ng/mL (R&D systems) and rhIL-15 at 1ng/mL (Miltenyi). Cytokines and medium were replaced every 2-3 days. Where indicated, increasing concentrations of DMSO (Sigma) were added from day 0, as given by % v/v. Cytokines, medium and DMSO were replaced at day 4/6/8/10.

### Human CD3^+^ T cell isolation and culture under DMSO exposure

Leukocyte-enriched buffy coat samples were obtained from healthy donors attending the Red Cross center after informed consent and ethical committee approval (EC2019-1083). Peripheral blood lymphocytes were prepared by a Ficoll-Paque density centrifugation as described in the instruction manual for Leucosep™ (Greiner bio-one). CD3^+^ T cells were isolated by negative selection (MojoSort™ Human CD3^+^ T cell selection kit, Biolegend) according to the manufacture’s protocol. CD8^+^ T cells were isolated from the same donor by negative selection (MojoSort™ Human CD8 T cell Isolation kit, Biolegend) according to the manufacture’s protocol. Cells were cultured in IMDM + Glutamax medium (Gibco-BRL) supplemented with 10% heat-inactivated fetal calf serum (FCS) and stimulated with Immunocult™ Human CD3/CD28 T cell Activator (Stemcell Technologies) (25 µL/ 10^6^ cells) for 3 days at 37°C in the presence of 10 ng/ mL IL-12 (Biolegend). Prior to cell seeding, cells were washed twice with PBS before putting them in culture with rhIL-7 at 10ng/mL (Miltenyi) and rhIL-15 at 10 ng/mL (Miltenyi). Cytokines, medium and DMSO were replaced every 2-3 days. Cell densities were maintained between 1 × 10^6^ and 3 × 10^6^ cells/ mL. Where indicated, increasing concentrations of DMSO (Sigma) were added from day 0 as given by % v/v.

### Proliferation assay

Human T cells were stained with Tag-it Violet™ Proliferation and Cell Tracking Dye (Biolegend), according to manufacturer’s protocol. Tag-it Violet™ passively diffuses into the cell where esterases cleave acetoxymethyl esters (AM) on the molecule, Tag-it Violet™ which then covalently attaches to intracellular proteins enabling its long-term retention.

### Flow cytometry analysis

For human T cell phenotyping, cells were labeled with fluorescent antibodies against human CD8, CD4, CD62L and CD45RA (BD Biosciences) and CD197 (CCR7), CD25, CD69, TIM-3 (CD366), CD152 (CTLA4) and CD279 (PD-1) (Biolegend). A Fixable dye eFluor™ 780 (eBioscience) was used to evaluate live/dead cells.

Data were acquired using a FACS Symphony™ equipped with five lasers (355, 405, 488, 561, 640 nm) (BD Biosciences) and analyzed with FlowJo Version 10 software (TreeStar). Flow cytometer calibration was performed using CS&T beads (BD Biosciences). The gating strategy was set based on fluorescence minus one (FMO) controls and retained for all samples. The gating strategy for the immunophenotyping of human CD8^+^ T cells is shown in Supplementary Figure 1.

### Statistical analyses

To test for the effect of DMSO on the fractions of T_N_, T_EFF_, T_EM_ and T_CM_ cells, we separately modeled the CD4^+^ cells in the CD3^+^ pool, the CD8^+^ cells in the CD3^+^ pool and the purified CD8^+^ cells (Figure 5) and IL-2 or IL-7/IL-15 stimulated cells (Figure 3). For each of these 5 settings, the outcome variable was the percentage of cells (belonging to a certain subtype) at day 13 (Figure 5) or day 11 (Figure 3) of DMSO stimulation respectively. We used a beta regression model with logit link function to model this outcome variable. As predictor variables we used the cell subtype, the % of DMSO and the T-cell donor, all as categorical variables. We allowed for two-way interactions of DMSO with subtype and donor with subtype, resulting in a final model: percentage ~ DMSO*subtype + donor*subtype. Furthermore, we allowed the dispersion to be dependent on the subtype as well. We fitted the models with the glmmTMB package^29^ in R^30^. Model quality was assessed with the DHARMa package^31^. Multiple testing corrected custom contrasts with their p-values and 95% confidence intervals were obtained using the multcomp package^32^. We contrasted each DMSO concentration with the control setting (no DMSO) for each T cell subtype.

## Acknowledgements

NF and LM are staff scientists of VIB. EDB was a predoctoral fellow at FWO during the project and had a doctor-assistant mandate at UGhent. EP is a research associate of VIB. This work was supported by grant G050420N of FWO Vlaanderen and by a Young Investigator Proof of Concept (YIPOC) grant of the Cancer Research Institute Ghent (CRIG). We are grateful to E. Wyseure for help with the T cell cultures. We thank the VIB Flow Core (https://vib.be/labs/vib-flow-core-ghent) facility for their services.

## Supplementary figures

**Supplementary Figure 1.**
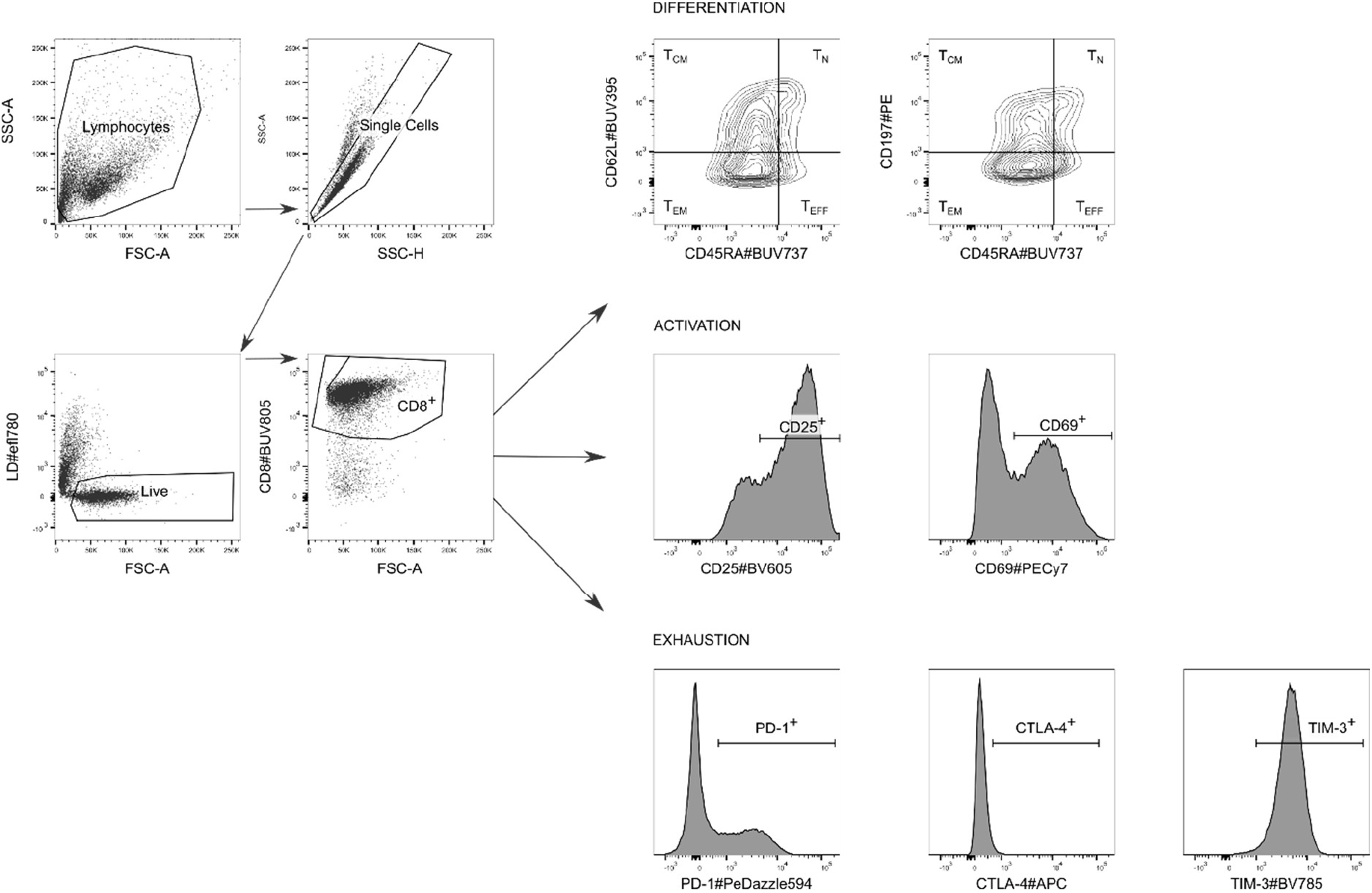
Gating strategy for the immunophenotyping of human CD8^+^ T cells. Total cells are first gated on the forward scatter (FSC)/ side scatter (SSC) plot to eliminate debris and single cells are then selected on the SSC-area (A)/ SSC-height (H) plot. CD8^+^ cells are subsequently gated on live cells. These are then further gated for the subsets of interest. Memory cells are separated according to CD45RA expression and then further phenotyped according to CD197 or CD62L expression as either T_N_ (CD45RA^+^CD197^+^ or CD45RA^+^CD62L^+^), T_EFF_ (CD45RA^+^CD197^-^ or CD45RA^+^CD62L^-^), T_EM_ (CD45RA^-^CD197^-^ or CD45RA^-^CD62L^-^) and T_CM_ (CD45RA^-^CD197^+^ or CD45RA^-^CD62L^+^). The activation state is evaluated based on the expression of CD25 and CD69, while T cell exhaustion is assessed based on the expression of PD-1, CTLA-4 and TIM-3.

**Supplementary Figure 2.**
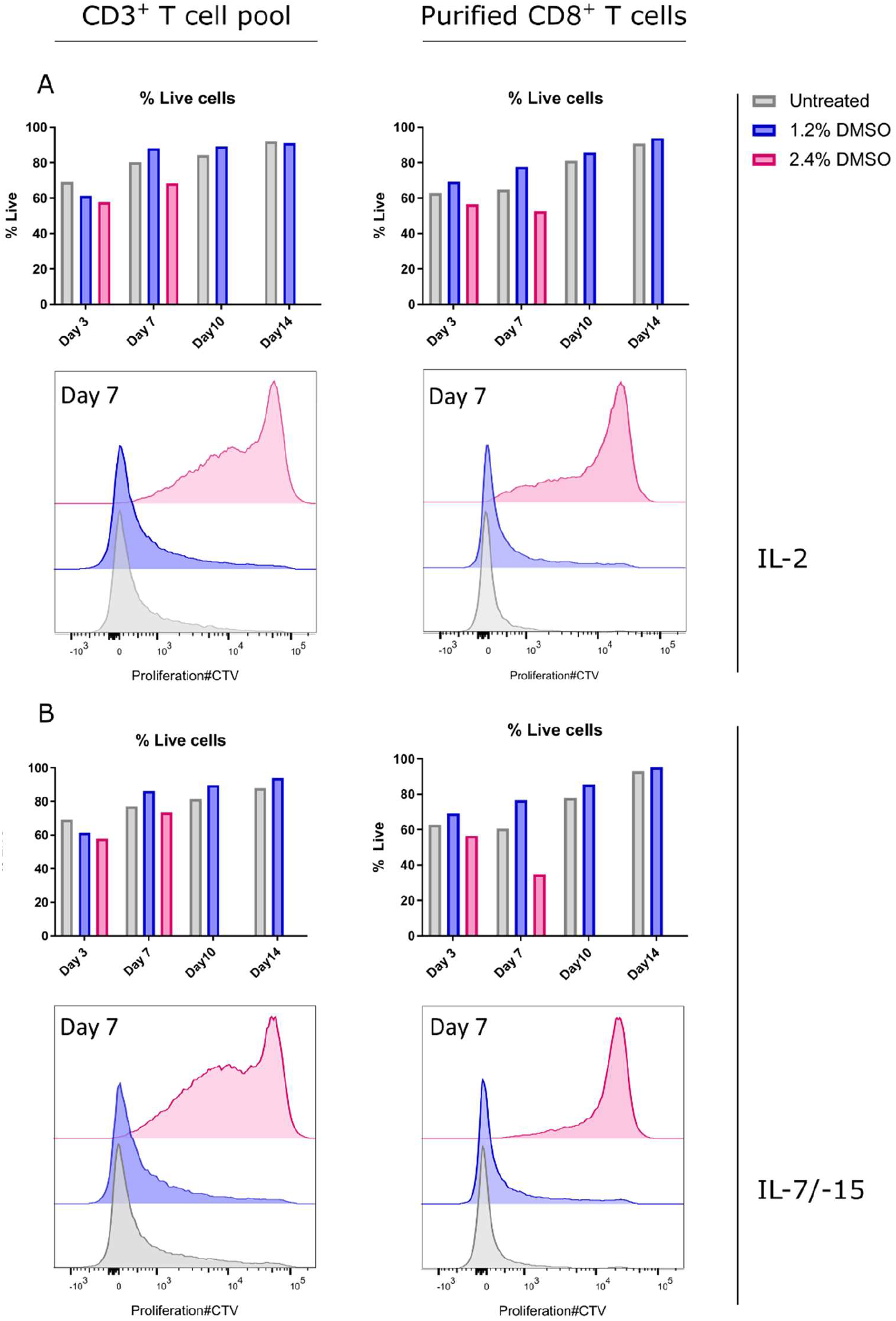
The effect of 2,4% DMSO concentration on human CD3^+^ and CD8^+^ T cell viability and proliferation. Purified human CD3^+^ T cells and CD8^+^ T cells (from the same donor) were stimulated for 3 days in the presence of IL-12 and cultured in the presence of IL-2 (A) or IL-7 and IL-15 (B) and two concentrations of DMSO (1.2%-purple, 2.4%-pink) or left untreated (grey). Viability and proliferation were analyzed by flow cytometry on different days as indicated on the X-axis, on either the CD3^+^ pool (left panels) or CD8^+^ purified T cells (right panels). On day 10 and day 14, no measurement was possible for the 2.4% DMSO condition since all cells died by that timepoint.

